# Millisecond dynamics of BTK reveal kinome-wide conformational plasticity within the apo kinase domain

**DOI:** 10.1101/135913

**Authors:** Mohammad M. Sultan, Rajiah Aldrin Denny, Ray Unwalla, Frank Lovering, Vijay S. Pande

## Abstract

Bruton tyrosine kinase (BTK) is a key enzyme in B-cell development whose improper regulation causes severe immunodeficiency diseases. Design of selective BTK therapeutics would benefit from improved, *in-silico* structural modeling of the kinase’s solution ensemble. However, this remains challenging due to the immense computational cost of sampling events on biological timescales. In this work, we combine multi-millisecond molecular dynamics (MD) simulations with Markov state models (MSMs) to report on the thermodynamics, kinetics, and accessible states of BTK’s kinase domain. Our conformational landscape links the active state to several inactive states, connected via a structurally diverse intermediate. Our calculations predict a kinome-wide conformational plasticity, and indicate the presence of several new potentially druggable BTK states. We further find that the population of these states and the kinetics of their inter-conversion are modulated by protonation of an aspartate residue, establishing the power of MD & MSMs in predicting effects of chemical perturbations.

## Introduction

Protein kinases are regulators of biochemical pathways in eukaryotic cells^1–3^ responsible for initializing and controlling signaling cascades^3,4^ by catalyzing the transfer of ATP’s gamma phosphate group to target residues on other enzymes. Given their critical involvement in cellular processes, their functionality is tightly controlled through a combination of regulatory domains^2,5,6^ and post-translational modificiations^3^ that modulate their multi-state behavior^1,6–9^. Sequence mutations, truncations, and over-expression of various kinases have been phenotypically linked to various cancers^3,4,10^ and other diseases^11^.

Bruton tyrosine kinase (BTK)^12–15^, part of the TEC family of kinases, is involved in T-cell and B-cell development. In humans, poor B-cell maturation leads to severe immune deficiencies, including increased susceptibility to bacterial infections^16^. Therefore, BTK’s catalytic domain is a pharmaceutical target with several inhibitors, including FDA-approved drugs^17^. Similar to other kinases, BTK’s catalytic domain (Figure 1a) is bi-lobal with a β-sheet heavy N-lobe and a α-helical C-lobe. ATP and magnesium ions bind in the active site between the two lobes (Figure 1a). The activation loop (A-loop, Figure 1a, red) connects the two lobes and modulates substrate binding. Phosphorylation of Tyr551 at the C-terminal end of the A-loop increases BTK’s activity by ten fold^15^. The N-terminal end of the A-loop contains the highly conserved aspartate-phenylalanine-glycine motif (DFG, Figure 1b, blue) which samples several pharmacologically relevant states^18^. The aspartate of the DFG can be protonated^9^, modulating drug binding^19^. In the N-lobe, the glycine rich phosphate-positioning loop (P-loop, Figure 1a, green) helps to position ATP’s gamma phosphate group. One β-sheet over, the N-lobe contains a conserved Lysine (Lys430, Figure 1b) that hydrogen bonds with ATP’s α-phosphate group. The N-lobe also has the catalytic helix (C-helix, Figure 1b, orange) which contains a conserved Glutamate residue (Glu445, Figure 1b) that forms critical salt bridges to Lys430 in the active state and Arg544 in the inactive form.

**Figure 1:**
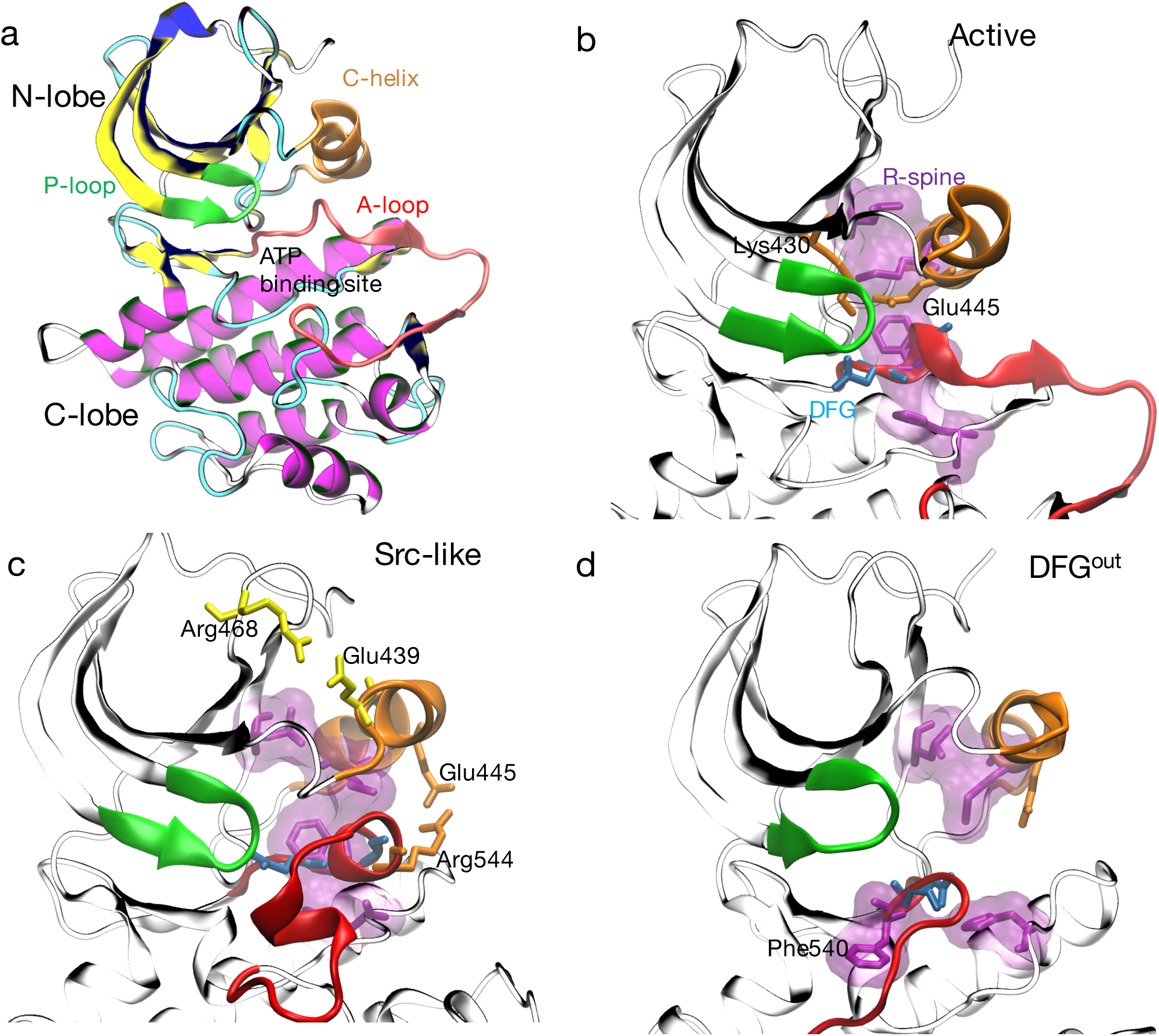
*BTK exists in several thermodynamically stable states. Within the MD ensemble, BTK catalytic domain samples several states, including (a) active (DFG*^*in*^*/C-helix*^*in*^*) (b), Src-like (A-loop folded/DFG*^*in*^*/C-helix*^*out*^*) (c) and DFG*^*out*^ *(d) within our simulation ensemble. The transition from active (a) to Src like (b) is defined by the outward rotation of the C-helix (orange) and folding of the A-loop (red). The C-helix rotation breaks a critical salt bridge between Glu445-Lys430 and forms salt bridges between Glu439-Arg468 (yellow) and Glu445-Arg544(orange). In the DFG*^*out*^ *state, Phe540(purple), part of the DFG motif (blue) rotates away from the core of the protein towards the ATP binding site. The R-spine (purple surface) forms continous hydorphobic contacts in the active state but is broken in the other states.*

Crystallographic and biochemical studies on BTK^13–15^ and other kinases^3,20^ have already provided a considerable amount of insight into their thermodynamically accessible enzymatic states. BTK can exist in active^14^, inactive^21^, and DFG out^21^ states. In the putative active state (Figure 1b), the DFG-aspartate residue moves towards the ATP binding site (DFG^in^) to chelate magnesium; the C-helix rotates into the protein core (C-helix^in^)^14^; and the A-loop is unfolded and transiently samples a β-sheet secondary structure. In the DFG^in^ inactive state, the C-helix rotates outwards (C-helix^out^), and the A-loop folds into a double helix (Figure 1c)^14^. We refer to this double helical inactive state as Src-like due to it topological similarity to the Src kinase’s inactive state^22^. In the DFG^out^ state, the DFG-Asp rotates towards the core of the protein (Figure 1d, Supporting Table 1).

While the crystallographic coordinates for BTK provide us with structural insights, these models do not provide information about unrealized thermodynamically stable states or the pathways connecting them. Molecular dynamics (MD) is a computational modeling technique used to complement experimental work in biophysical systems. MD^7,9^ provides atomistic insight into complex processes, and has led to the proposal of several new kinase structural intermediates^7,23–25^ and allosteric pathways^8^.

In this paper, we performed an aggregate of 1.7 milliseconds of MD simulations on the DFG-deprotonated (BTK-ASP) and DFG-protonated (BTK-ASH) forms of the unliganded BTK catalytic domain on the massively distributed Folding@home^26^ computing platform(see Methods for details regarding the homology modeling). The aggregate simulation times make this study three times longer than the largest reported MD results on kinases^7^ and three orders of magnitude larger than any computational investigation into BTK’s dynamics^12,27^. We characterized kinome-wide structural plasticity within the C-helix and DFG motifs, identifying a number of conformations as viable pharmaceutical targets. We used Markov state models^28,29^ (MSMs) to gain atomistic insight into the thermodynamics and kinetics of BTK’s conformational ensemble, identifying a structurally diverse intermediate state that links the active, Src-like, and DFG^out^ states.

### BTK kinase domain samples kinome-wide conformational space

We began our analysis by comparing the structural heterogeneity in the MD BTK-ASP dataset to the kinome-wide PDB classification of Möbitz et al^18^ (Figure 2). In that paper, the authors classified kinase structures along variations of the conserved DFG motif, the C-helix, and the A-loop. Starting from several publically available BTK protein coordinates (Supporting Table 2), our simulations capture kinome-wide crystallographically observed states that were previously stabilized via a combination of sequence, drugs, small peptides, and crystallographic conditions. For example, our simulations predict several configurations that Möbitz classified as the Imatinib, a leukemia drug, binding mode (Supporting Figure 3) for Abl kinase^19,30,31^. In this pose, the DFG-Phe540 residue (Figure 1d) rotates towards the ATP binding site (Figure 1a), creating a back pocket capable of accepting an aromatic moiety. It is worth emphasizing that the A-loop (including DFG-Phe540) was not resolved in similar BTK^21^ structures, creating difficulties for structure-based drug discovery. This result indicates the increasing ability of MD simulations to predict physiologically and pharmacologically relevant positioning of critical structural motifs. Intriguingly, BTK’s plasticity supports the model that all kinases sample a single conformational landscape whose topology is modulated by their sequence and/or chemical environment.

**Figure 2:**
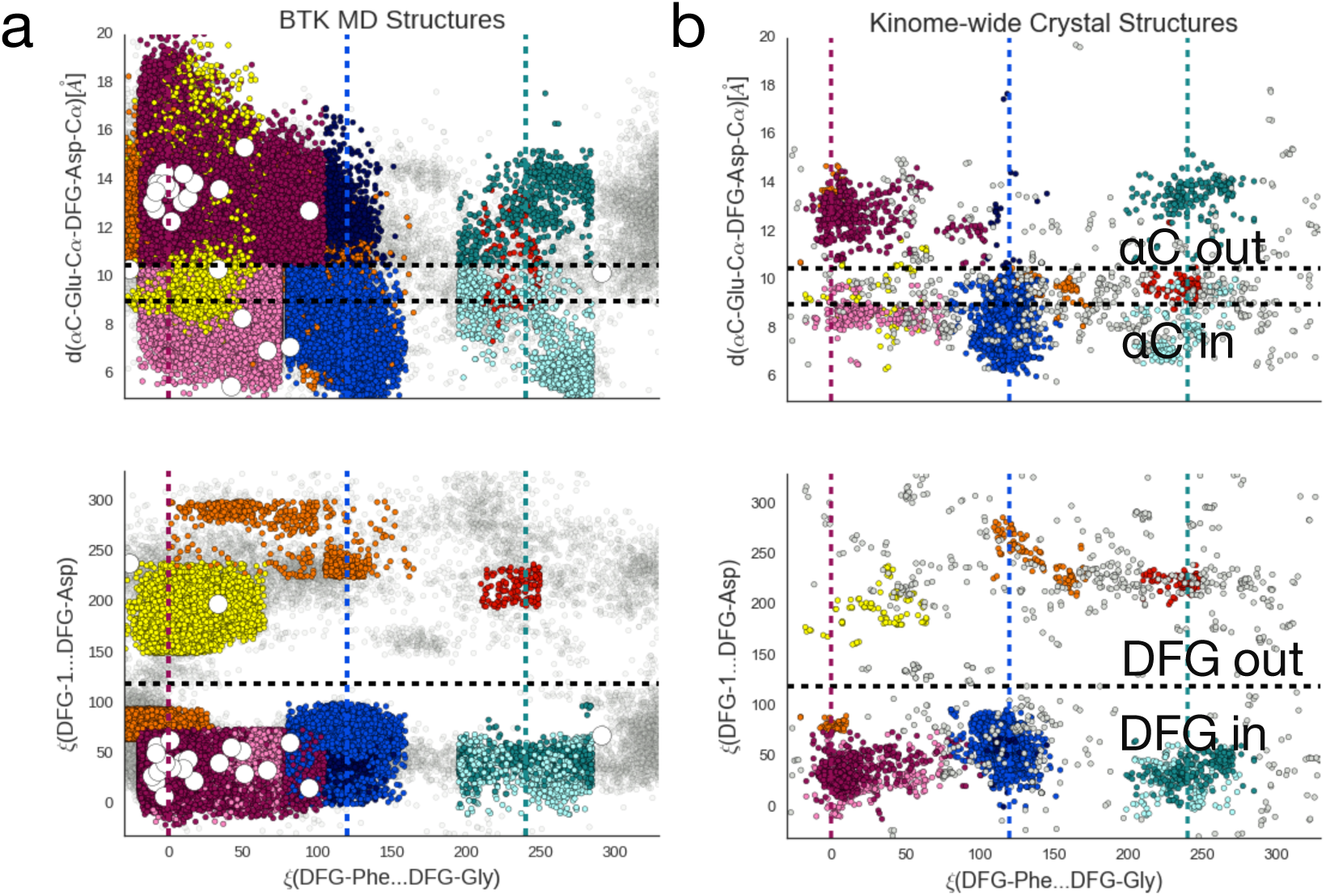
*BTK’s apo domain contains kinome-wide conformational plasticity. Comparison of 9% of MD generated structures (a) against publically available kinase domain structures (b) projected along three key degrees of freedom as outlined in Möbitz et al*^18^. *We used the data and classification scheme provided in ref.* 18 *to generate (b). The top y-coordinate tracks the C-helix*^*in*^ *to C-helix*^*out*^ *transition while the bottom y-coordinate tracks the DFG*^*in*^ *to DFG*^*out*^ *transition. The common x-axis subdivides the conformations into pharmacologically relevant states of the DFG motif. The white circles in (a) correspond to the starting configurations for the MD simulations. The points are colored according to their Möbitz classification and detailed in Supporting Figure 16. For BTK’s free energies along these coordinates, see Supporting Figure 9.*

### BTK’s apo domain is primarily inactive

To gain insight into the thermodynamics and kinetics of BTK, we built a statistically robust MSM for the hundreds of collected MD trajectories (Supporting Figure 1-2). Markov modeling^28^ involves Voronoi partitioning of the accessible phase space into states and counting the transitions between the states. The metastable states are defined using a kinetically relevant distance metric (see Methods) that is learnt via sparse time structure-based independent component analysis (sparse-tICA)^32–35^. tICA finds linear combinations of input MD features that de-correlate the slowest within the given dataset. The dominant components – tICs – relate the slow structural changes to long timescale protein dynamics. After performing this dimensionality reduction, we built an optimized^36,37^ MSM (see Methods) whose transition matrix reflects the ensemble equilibrium populations and long timescale processes.

We projected the BTK-ASP and BTK-ASH MD ensembles onto the dominant tICs (Figure 3a) and several structural order parameters (Supporting Figure 9-11). These structural projections help us to understand the relative thermodynamic stability of various states as a function of the chosen order parameter. Our analysis indicates that the two principal tICs correspond to (1) the flipping of the DFG motif (Supporting Figure 6) and (2) the unfolding of the A-loop coupled with the rotation of the C-helix to the protein core, respectively (Supporting Figure 7). The free energy surface (Figure 3a) lets us define a four-state model (Figure 3b, Supporting Figure 8) in which a structurally heterogeneous intermediate hub (I_1_, Supporting Figure 5) controls access to active, Src-like, and DFG^out^ states. This intermediate contains a partially or fully unfolded A-loop, DFG^in^, and C-helix^out^. Our simulations predict the Src-like state to be the most stable state of BTK-ASP (Figure 3a) with the active and DFG^out^ states both being 1-2 kcal/mol above the minimum energy observed. Within the BTK-ASP model, these states (Supporting Figure 8, 10 & 12) have populations of 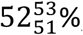, 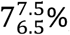, and 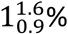 respectively. The sub-script and super-script indicate the 95% confidence interval. See Supporting Figure 2 for the complete distribution. The rest of the population exists in the intermediate state. The low active state population lines up with the large number of inactive crystal structures (Figure 2a, dark magenta region), and biochemical studies showing that BTK’s Tyr551’s phosphorylation^15^ is required for full activation. The number of minor and major populated states, and their micro to millisecond exchange timescales, are also consistent with NMR studies on other kinases. For example, the DFG-Phe in apo-p38α^38^ is unobserved in NMR due to line broadening, attributed to DFG-flip conformational exchange. Similarly, dual phosphorylation on ERK2^39^ shifts the equilibrium to its active state by about 3kcal/mol while ligand binding to protein kinase A^38^ induces slow inter-domain motion. This suggests that the solvated kinase catalytic domain samples a diffuse free energy landscape whose topology can be modulated by a number of factors. MSMs present a natural framework to handle these perturbations, and we next focus on the effects of one of them namely protonation of the DFG-Asp539.

**Figure 3:**
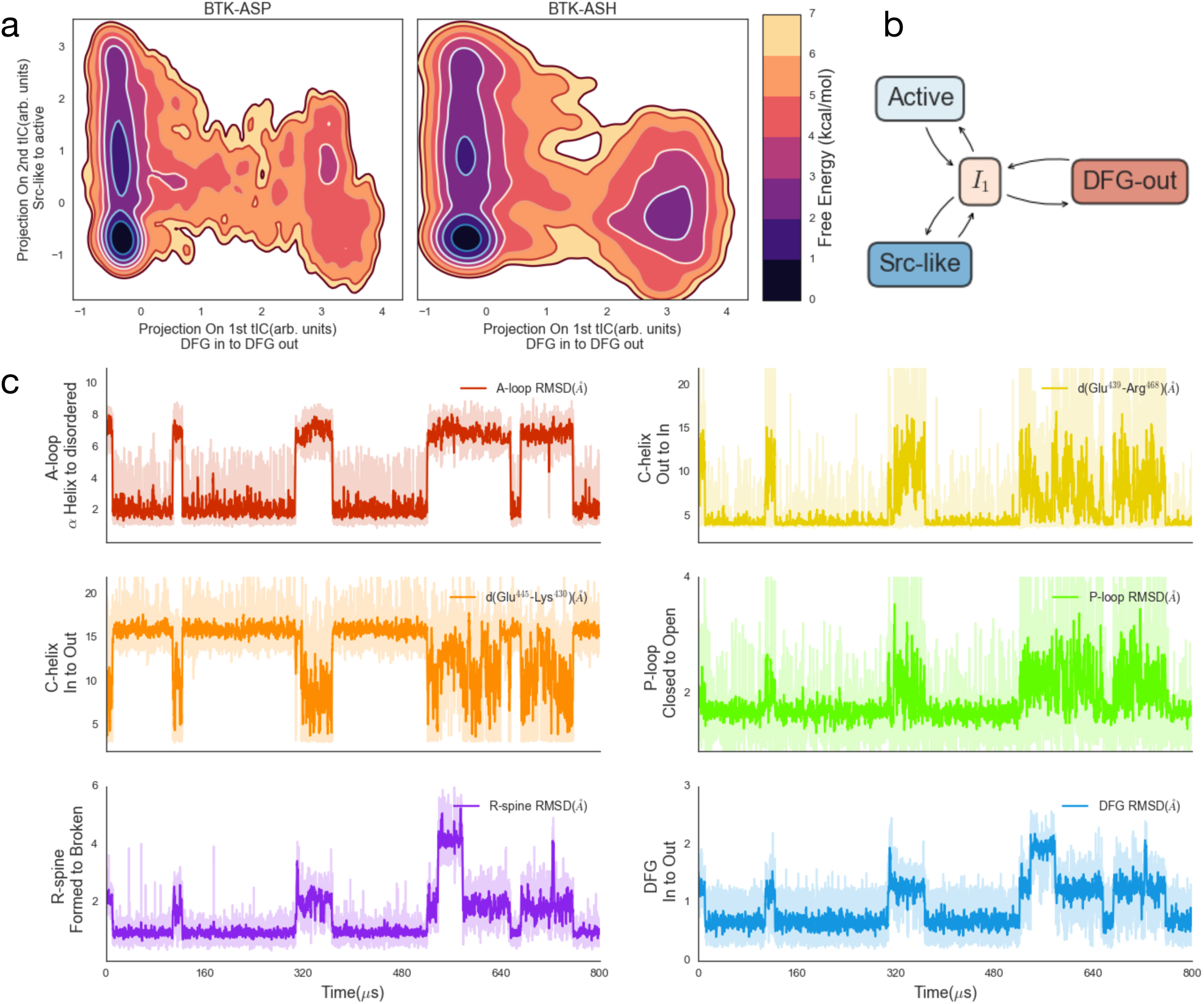
*MSMs predict a multistate ensemble whose populations are modulated via DFG protonation. Thermodynamics of the BTK-ASP and BTK-ASH ensembles projected along the two dominant tICs (a) show a stable Src-like state. For standard errors along each coordinate, see Supporting Figure 12. Simple four state cartoon model (b) of the kinase dynamics. Kinetics of several molecular switches as a function of time along a MSM trajectory for BTK-ASP (c). The MSM trajectory was generated using a Monte Carlo algorithm to simulate a trajectory of 800 μs from the Markovian transition matrix. At each step, we randomly selected a simulation structure assigned to that state to report the instantaneous observables. The root mean squared deviation (RMSD) of the A-loop is calculated using the heavy atoms of residue Asp539-Phe559. For A-loop RMSD to the extended state, see Supporting Figure 17. We used the delta carbon of the Glu439 and zeta carbon of Arg468, and the delta carbon of Glu445 and zeta nitrogen of catalytic Lys430 to calculate the distances in the next two panels to quantify C-helix in to out transition. We used Thr410-Val415, and Phe540, Met449, His519, and Leu460 heavy atoms to quantify the P-loop, and R-spine RMSD. The R-spine is only completely formed when the C-helix (orange trace) is rotated inwards. The DFG RMSD is calculated using heavy atoms from Asp539-Gly541. For all RMSD calculations, we used a double helical inactive state as the reference state. The lighter color traces give the instantaneous value for the observable and the dark traces provide moving averages across 10 frames. The color corresponds to the color scheme used in Figure 1 to highlight structural motifs in BTK.*

Based upon pKa calculations^40^(Supporting Note 2), and past MD studies on EGFR, Src, and Abl kinases^9,19,41^, we next selectively protonated DFG-Asp539 (BTK-ASH) to quantify its effects on thermodynamics and kinetics. Our models predict that the DFG^out^ state is stabilized by approximately 1 kcal/mol (relative to BTK-ASP’s DFG^out^) upon protonating the aspartate (Figure 3a). Compared to BTK-ASP, the increase in DFG^out^ population comes from the reduced free energy cost^19^ of putting a neutral protonated DFG-Asp539 in a hydrophobic environment (Figure 1d, Supporting Figure 19). Within the BTK-ASH simulation set, we find that the Src-like, active, and DFG^out^ states have populations of 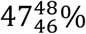, 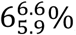, and 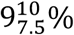 respectively. Furthermore, we observed that protonation accelerates the DFG flip. To quantify this effect, we calculated the median value for the mean first passage time (MFPT)^42,43^ between DFG^in^ and DFG^out^ states (Supporting Figure 2e and Supporting Figure 8). Starting from the DFG^in^ states, the median value for the MFPT to the DFG^out^ state reduces from 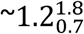 in BTK-ASP ensemble to 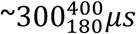 in the protonated BTK-ASH ensemble. The reverse value remains relatively similar 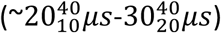. See SI Figure 2e and 2f for the full distribution of macro state populations and median DFG MPFTs across several hundred rounds of bootstrapping.

To understand BTK-ASP’s kinetics, we sampled multiple long trajectories from the BTK-ASP microstate Markovian transition matrix using a kinetic Monte Carlo algorithm (see Methods). The stochastic algorithm allows us to stitch the shorter trajectories (hundreds of nanoseconds) into a longer “mock” trajectory that would be otherwise inaccessible by traditional sampling. A representative 800 *μs* trajectory projected along several of the dynamical regions within the kinase domain is illustrated in Figure 3c. The model predicts that the A-loop is quite flexible, sampling conformations spanning RMSDs on the order of 10s of angstroms (Supporting Figure 17). Such structural heterogeneity within the A-loop has been widely reported in kinase crystal structures^6^ and MD models of apo^9^ and ATP^7^ bound kinases. We find that these transitions are coupled to the outward rotation of the C-helix and flipping of the DFG motif. The outward transition of the C-helix also breaks the regulatory spine (R-spine)^1,44^, which consists of the four conserved residues Met449 (part of the C-helix), Leu460, Phe540 (part of conserved DFG motif) and His519 (part of the conserved HRD motif). Over the course of the simulation, the R-spine samples three distinct macrostates (Figure 3c purple trace) corresponding to the Src-like, active, and DFG^out^ states. In the active state, the R-spine forms a rigid, continuous hydrophobic surface stabilized via multiple Van der Walls interactions. These interactions are broken in the inactive states (Supporting Table 1) where the Met449 moves out of the core of the protein and Phe540 adopts a range of configurations (Figure 1).

Interestingly, our model predicts that the kinase deactivation to a Src-like state (A-loop folded, DFG^in^, & C-helix^out^, Figure 3c between the 350-450*μs* mark) quenches motions in the dynamic glycine-rich phosphate positioning loop (P-loop, Figure 3c green trace). The P-loop samples open and closed configurations in the active and DFG^out^ states. However, the P-loop closes when the A-loop folds. Our model indicates this rigidity is due to both the formation of a new backbone hydrogen bond (Supporting Figure 14) between Lys433 and Phe413 and favorable p-stacking (Supporting Figure 14) between Leu542 and Phe413.

### Deactivation proceeds through an intermediate state

We now turn to the structural changes involved in the two dominant apo BTK transitions, namely active (C-helix^in^/DFG^in^) to Src-like (A-loop folded/C-helix^out^/DFG^in^) and DFG^out^ to DFG^in^. Like most kinases^44–46^, BTK’s active state features the C-helix rotated towards the core of the protein, enabling the formation of a critical^7,47^ salt bridge between the Glu445 and Lys430 (Figure 4a). The A-loop is extended and transiently forms a beta-sheet.

**Figure 4:**
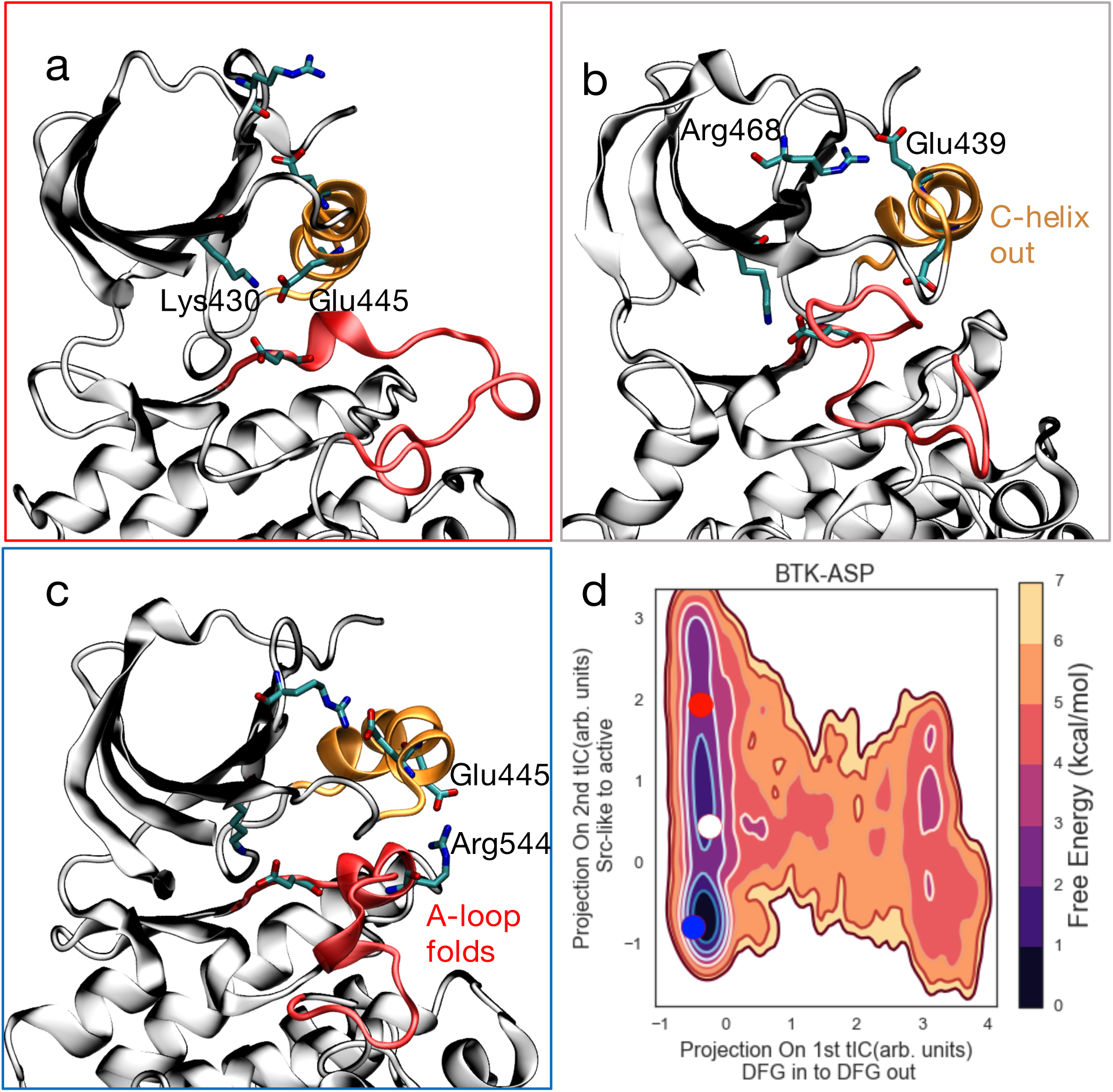
*BTK’s deactivation proceeds via an intermediate state. Starting from the active state (a), the C-helix swings out to form an intermediate (b) characterized by a disordered A-loop and a stable Arg468-Glu439 salt bridge. The activation loop then folds into a Src-like double helical inactive state (c). The double helical state is stabilized by a secondary salt bridge between the catalytic Glu445 and Arg544. The P-loop has been omitted in all three panels for the sake of clarity. The heat map (d) shows the projection of the centroids of these states unto our free energy landscape. Panel (d) has been reproduced from Figure 3 for clarity.*

Deactivation to a Src-like state exposes an allosteric binding pocket. In both the BTK-ASP and BTK-ASH ensembles, deactivation to a Src-like state follows a two-step^7,9,19,24,25^ process. In the first step, the C-helix rotates out of the core of the protein (Supporting Figure 13) to a metastable intermediate conformation (Figure 4b). In several of our trajectories the C-helix rotation was preceded by a backbone shift at the N-terminus region of the A-loop. This intermediate is stabilized by a salt bridge between Arg468 and Glu439. The A-loop is still relatively unstructured but transiently samples partially helical states (Supporting Figure 5). The outward rotation of the C-helix to the catalytically inactive intermediate state opens an allosteric pocket (Supporting Figure 4)^7^, which can potentially be used to design selective BTK inhibitors. Supporting Movie 1 contains an example of one of our BTK-ASP trajectories that spontaneously goes from the active to the intermediate.

Starting from the intermediate state, the A-loop folds into a double helix. This state forms a deep free energy basin in our MSM (Figures 4c-d). The inactive state is stabilized by the presence of two complementary salt bridges (Glu439-Arg468 and Glu445-Arg544^5,7,8^, Figure 4c). Within several milliseconds of aggregate sampling, we do not observe a deactivation event where the A-loop folds prior to the outward rotation of the C-helix, suggesting that this pathway is thermodynamically inaccessible.

### DFG flip occurs via the C-lobe

We observed several partial and complete DFG transitions for the BTK-ASP and BTK-ASH ensembles in trajectories that lasted hundreds of nanoseconds to a few microseconds (Supporting Movies 2-4). For both ensembles, our MSMs inferred the equilibrium populations and kinetics using all of the trajectories, regardless of whether they contained a crossover event (Figure 3 and Figure 5d).

**Figure 5:**
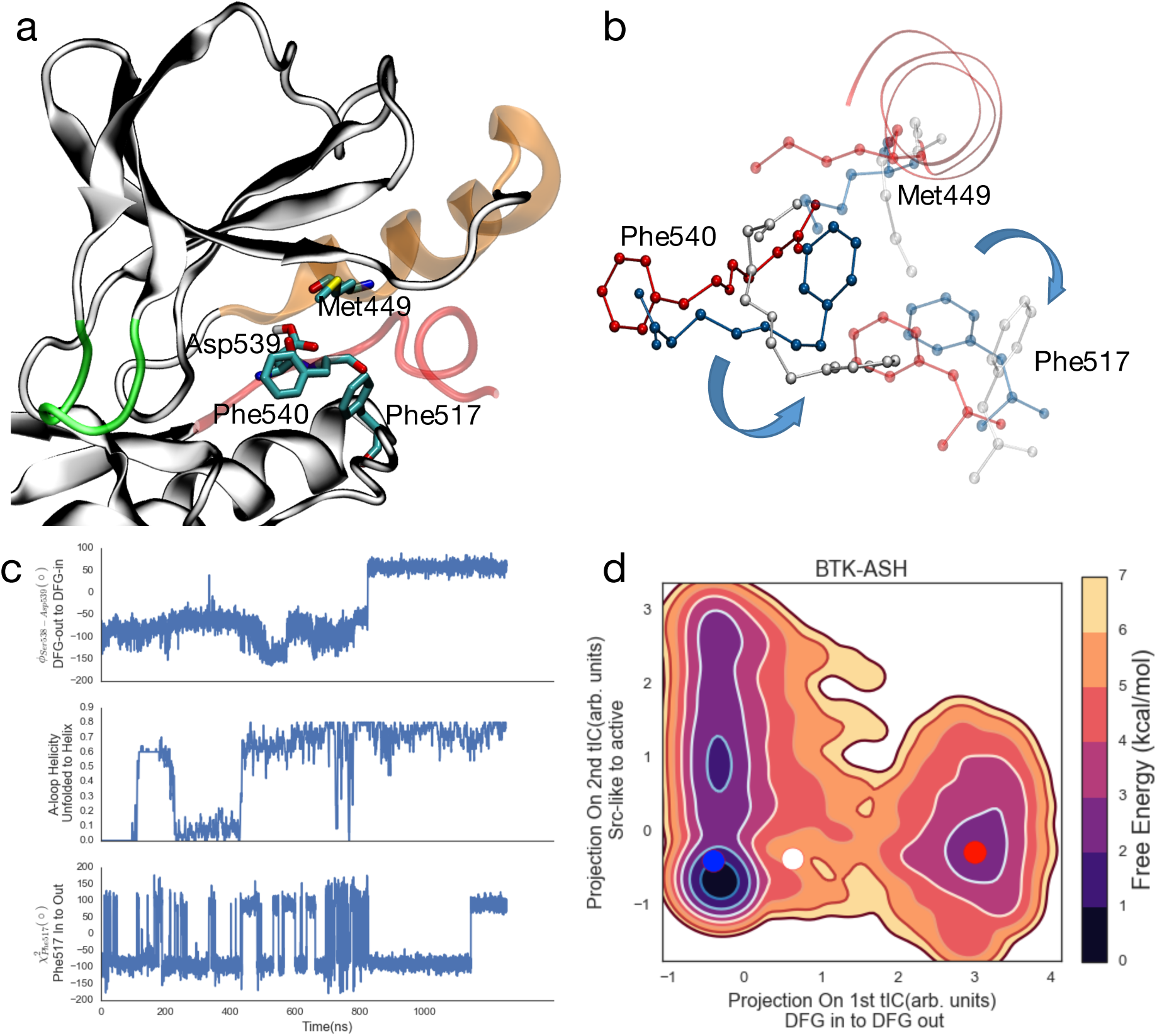
*BTK DFG flips via the C-lobe, and proceeds after the formation of a helical intermediate state (a). Snapshots (b) going from red to white to blue, from the DFG*^*out*^ *to DFG*^*in*^ *trajectory showing the transient outward rotation of Met449 and Phe517 for the DFG flip. The DFG*^*out*^ *to DFG*^*in*^ *cross-over (c, panel 1) is preceded by the folding of residues Ser543-Leu547 (c, panel 2) and transient outward rotation of both Met449 and Phe517 (c, panel 3). Projection of the 3 selected frames from (b) onto the top two tICs (d) gives us the approximate free energies of the DFG*^*out*^, *intermediate and DFG*^*in*^ *states.*

Figure 5 shows the details of one of the transition trajectories from the BTK-ASH ensemble starting from the 3OCT crystal structure with an unfolded A-loop and DFG^out^ state (Supporting Movie 2). Within the trajectory, the A-loop first folds into the ATP binding site, forming a helical intermediate (Figure 5c, Supporting Figure 18). In this intermediate, residues Leu542 to Leu547 form a helical turn that folds into the kinase’s core though the rest of the A-loop remains mobile. This intermediate has been previously reported^9,19^ for the EGFR kinase. Furthermore, Kuglstatter et al^21^ reported a DFG^in^ crystal structure for BTK where the the A-loop folds into the ATP binding site, demonstrating its [meta]stability. Within our simulation, the BTK-ASH molecule samples this intermediate until the DFG-Phe rotates towards the core of the protein (Figure 5b, white) coming on to the same side as the DFG-Asp. In the last step, the DFG-Asp moves into the ATP binding site, completing the crossover transition.

While no unbiased MD simulation results exist for the DFG flip for either BTK-ASP or BTK-ASH molecule, the pathway presented here diverges from EGFR kinase’s DFG-flip pathway^9^ in two aspects. In the previous simulations, the DFG-Phe residue flips by moving across the N-lobe of the kinase. Within our model, the phenylalanine residue exclusively moves via the C-lobe of the kinase while the aspartate moves via the N-lobe (Supporting Figure 15). This sequence is conserved in both the full and partial trajectories (Supporting Movies 2-4) that transition from the DFG^in^ state to the DFG^out^ state and vice versa. To our knowledge, this mechanism has never been proposed, though that is likely due to the computational difficulties of simulating the DFG flip transition. It is worth noting that our distributed computing approach^26^ allowed us to sample the DFG flip in an unbiased fashion using commodity GPUs, and our MSM was able to capture the DFG transition as the slowest mode within our tICA model. Secondly, while we do observe the presence of a helical intermediate state in several of the transitions, we also observed a DFG^out^ to DFG^in^ transition for the BTK-ASP molecule in which the A-loop remains unstructured (Supporting Movie 3). The litany of flipping pathways follows from the inherent stochasticity of molecular conformational change and emphasizes the need for extensive sampling and robust statistical modeling.

Three residues sterically and chemically hinder the DFG transition. The conserved catalytic Lys430 hydrogen bonds with the DFG-Asp539 while the conserved Met449 sterically prevents rotation of the DFG-Asp539 towards the ATP binding site. On the other side, Phe517 (Figure 5b-c) hinders the rotation of the DFG-Phe540 towards the protein core. Previous MD studies^9,19^ of the DFG flip observed spontaneous DFG transitions upon *in-silico* mutations of Met449, or its equivalent residue, and protonation of the DFG-Asp. As previously proposed, our data supports that protonation of the DFG-Asp can increase the likelihood of a spontaneous DFG flip. Our comparison of BTK-ASH to BTK-ASP showed that the DFG^out^ state is stabilized by 1kcal/mol upon DFG-Asp539 protonation. However, the combined effect of both DFG protonation and Met449 mutation remains to be observed. Lastly, while our simulations showed that the collapse of the folded A-loop into the kinase core predominantly precedes the DFG flip, it is not necessarily required, highlighting the ensemble nature of the pathway.

To summarize, the present results offer a detailed atomistic description of the thermodynamics and kinetics of the protonated and deprotonated forms of the apo BTK catalytic domain. Our model predicts that the apo kinase domain samples a range of conformational states that are yet to be crystallized but for which equivalent structures from other kinase domains exist. We complete structural modeling of a DFG^out^ binding pocket for BTK, which could potentially be used to design a new class of BTK inhibitors. Furthermore, our model indicates that a structurally diverse intermediate state connects the active, Src-like, and DFG^out^ states. For the first time, our results provide estimates for the equilibrium populations of all three dominant kinase states within a single model.

While we have chosen to separately analyze the BTK-ASP and BTK-ASH ensembles, the BTK ensemble in solution is a combination of both modulated by the DFG-Aspartate’s pKa in each microstate. Modeling this ensemble coupling would ideally require the use of constant pH simulations but can also be done post-hoc by analytical mixing of the parameterized transition matrices. This would entail running short constant pH(or QM/MM^48,49^) simulations to link the microstates, and is the subject of future research. Perhaps more interestingly, our BTK-ASP and BTK-ASH MSMs use a single set of state definitions, allowing us to explicitly compare relative free energies of differing kinase states. This can be extended to understanding the thermodynamic and kinetic effects of small molecule binding, mutations, regulatory domains, and post-transitional modifications. Given enough computational resources, it is theoretically possible to model all 602 known BTK mutations in the X-linked agammaglobulinemia database^16^. Such detailed atomistic characterization could be used to design the next generation of personalized and specific kinase inhibitors while increasing our understanding of the fundamental interplay between sequence and function.

## Methods

### Simulation setup

The Supporting information contains a more detailed methods section. Briefly, we downloaded 23 publically available BTK pdbs from the protein databank^50^, and used Modeller^51^ with default parameters to mutate out all the sequences to the human sequence for both BTK-ASP and BTK-ASH ensembles. We only kept the protein coordinates and removed all ligands (21 of the 23 structures had a co-crystal ligand). In cases, where the P-loop, C-helix or A-loop was unresolved, we modeled them in as an extended chain. Amber tools suite^52–54^ was used to solvate the protein structures in a water box and add counter ions. The Amber99sb-ildn^55^ force field was used to model protein dynamics in conjunction with the TIP3P^56^ water model. The structures were minimized in two steps using Amber and then loaded into OpenMM^57^ for NPT production runs on Folding@home^26^. Overall we generated 1.7ms of aggregate data for both ensembles.

### Markov state model

Building a MSM requires identification of metastable kinetically similar states. This splitting of the phase space is followed by counting the transitions between those states as observed in our trajectories at a Markovian (memory free) lag time. This transition model can be summarized using the following equation:

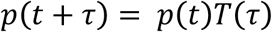

where *p*(*t*) is the probability distribution at time “t” while *p*(*t* + *τ*) is the probability distribution after a Markovian lagtime *τ*. Spectral decomposition of the MSM transition matrix was used to estimate the equilibrium populations and dynamical processes connecting those Markov states. The relaxation timescales for these dynamical processes can be obtained by using the following transformation on the associated eigenvalue *μ*

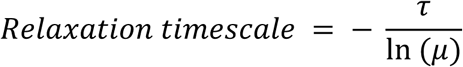

After sampling the MD trajectories using Folding@home, a total of 2,140 trajectories were vectorized using the protein dihedrals and selective closest heavy atom distances. This feature selection led to each frame being represented as a feature vector of length 5,532. We normalized the data and reduced its dimensionality using time structure independent component analysis (tICA)^32,35^. tICA seeks to find a set of linear combinations of features that de-correlate the slowest (at a certain lag time) while minimizing their correlation. This is done by solving the following generalized eigenvalue problem:

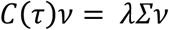

where *Σ* is the covariance matrix

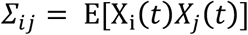

and *C*(τ) is the time lagged correlation matrix whose ij element is defined as

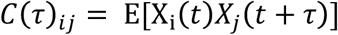

Here, *τ* is the tICA lagtime and can be different from Markovian lagtime. The tICA-transformed dataset was clustered using the K-means algorithm. We then used the cluster labeled dataset to build a MSM. For all projections, the deprotonated (BTK_ASP_) ensemble’s highest populated state was assigned an absolute free energy of 0 kcal/mol and all other free energies were reported relative to that state. Based upon previous work^7,58^ and the convergence of the implied timescales plot (Supporting Figure 1) for 50-500 state models, we chose a Markovian lag time of 80 ns. For all the other hyper-parameters, including the tICA lagtime, choice of kinetic mapping, number of tICA components, and number of cluster states, within the model, we turned to cross validation^36,37^. The parameters for the best model are given below:

| Hyper parameter | Value in best model |
| --- | --- |
| tICA lagtime | 208 ns |
| tICA number of components | 3 |
| tICA kinetic mapping | True |
| Number of clusters | 190 |

After we determined the optimal model given the current amount of sampling, we retrained the model on the entire set of trajectories. For the reported tICA model, we used a sparse variant of tICA^34^ for increased interpretability (Supporting Figure 6-7). The Markov transition matrix was fit via maximum likelihood estimation (MLE) with reversibility and ergodicity constraints. To obtain error bars for the equilibrium populations, 200 rounds of bootstrapping were performed over the original set of trajectories. The models were primarily analyzed using techniques laid out in previous papers^35,59^. To further query the model, we sampled an 800 μs long kinetic Monte Carlo trajectory (10,000 frames at a lagtime of 80 ns) from the Markovian transition matrix.

The trajectories were featurized and analyzed using the MDTraj^60^ package while tICA dimensionality reduction and Markov modeling were performed using MSMBuilder^61^. Most of the analysis was performed within the IPython/Jupyter scientific environment^62^ with extensive use of the matplotlib^63^, and scikit-learn libraries^64^. All protein images were generated using visual molecular dynamics (VMD)^65^, all protein surfaces were rendered using SURF^66^,and secondary structure was assigned using STRIDE^67^ as implemented in VMD.

## Acknowledgements

M.M.S acknowledges support from NSF-MCB-0954714. All authors acknowledge monetary support from the Pfizer Science & Technology budget. M.M.S would also like to thank Carlos Xavier Hernandez, Brooke Husic, Evan Feinberg, Ariana Peck, and other members of the Pande group for critical discussions and reading of the manuscript. The authors would like to thank Yatish Jain assisting with order parameter scripting and Mark Bunnage and John Mathias for their constant encouragement and support.

## References

1. Taylor, S. S. & Kornev, A. P. Protein Kinases: Evolution of Dynamic Regulatory Proteins. Trends Biochem. Sci. 36, 65–77 (2011).

2. Parsons, S. J. & Parsons, J. T. Src family kinases, key regulators of signal transduction. Oncogene 23, 7906–9 (2004).

3. Endicott, J. A., Noble, M. E. M. & Johnson, L. N. The structural basis for control of eukaryotic protein kinases. Annu. Rev. Biochem. 81, 587–613 (2012).

4. Hanks, S. K. & Hunter, T. The eukaryotic protein kinase superfamily: kinase (catalytic) domain structure and classification. FASEB J. 9, 576–96 (1995).

5. Levinson, N. M. et al. A Src-like inactive conformation in the abl tyrosine kinase domain. PLoS Biol. 4, e144 (2006).

6. Huse, M. & Kuriyan, J. The Conformational Plasticity of Protein Kinases. 109, 275–282 (2002).

7. Shukla, D., Meng, Y., Roux, B. & Pande, V. S. Activation pathway of Src kinase reveals intermediate states as targets for drug design. Nat. Commun. 5, 3397 (2014).

8. Foda, Z. H., Shan, Y., Kim, E. T., Shaw, D. E. & Seeliger, M. A. A dynamically coupled allosteric network underlies binding cooperativity in Src kinase. Nat. Commun. 6, 5939 (2015).

9. Shan, Y., Arkhipov, A., Kim, E. T., Pan, A. C. & Shaw, D. E. Transitions to catalytically inactive conformations in EGFR kinase. Proc. Natl. Acad. Sci. U. S. A. 110, 7270–5 (2013).

10. Greuber, E. K., Smith-Pearson, P., Wang, J. & Pendergast, A. M. Role of ABL family kinases in cancer: from leukaemia to solid tumours. Nat. Rev. Cancer 13, 559–71 (2013).

11. Jope, R. S. & Johnson, G. V. W. The glamour and gloom of glycogen synthase kinase-3. Trends Biochem. Sci. 29, 95–102 (2004).

12. Wang, Q. et al. Autoinhibition of Bruton’s tyrosine kinase (Btk) and activation by soluble inositol hexakisphosphate. Elife 4, e06074 (2015).

13. Mohamed, A. J. et al. Bruton’s tyrosine kinase (Btk): function, regulation, and transformation with special emphasis on the PH domain. Immunol. Rev. 228, 58–73 (2009).

14. Marcotte, D. J. et al. Structures of human Bruton’s tyrosine kinase in active and inactive conformations suggest a mechanism of activation for TEC family kinases. Protein Sci. 19, 429–39 (2010).

15. Dinh, M. et al. Activation mechanism and steady state kinetics of Bruton’s tyrosine kinase. J. Biol. Chem. 282, 8768–76 (2007).

16. Väliaho, J., Smith, C. I. E. & Vihinen, M. BTKbase: the mutation database for X-linked agammaglobulinemia. Hum. Mutat. 27, 1209–17 (2006).

17. Hendriks, R. W., Yuvaraj, S. & Kil, L. P. Targeting Bruton’s tyrosine kinase in B cell malignancies. Nat. Rev. Cancer 14, 219–232 (2014).

18. Möbitz, H. The ABC of protein kinase conformations. Biochim. Biophys. Acta - Proteins Proteomics 1854, 1555–1566 (2015).

19. Shan, Y. et al. A conserved protonation-dependent switch controls drug binding in the Abl kinase. Proc. Natl. Acad. Sci. U. S. A. 106, 139–44 (2009).

20. Sicheri, F. & Kuriyan, J. Structures of Src-family tyrosine kinases. Curr. Opin. Struct. Biol. 7, 777–85 (1997).

21. Kuglstatter, A. et al. Insights into the conformational flexibility of Bruton’s tyrosine kinase from multiple ligand complex structures. Protein Sci. 20, 428–436 (2011).

22. Xu, W., Doshi, A., Lei, M., Eck, M. J. & Harrison, S. C. Crystal structures of c-Src reveal features of its autoinhibitory mechanism. Mol. Cell 3, 629–38 (1999).

23. Ozkirimli, E., Yadav, S., Miller, W. & Post, C. An electrostatic network and long-range regulation of Src kinases. Protein Sci. 295, 1871–1880 (2008).

24. Gan, W., Yang, S. & Roux, B. Atomistic view of the conformational activation of Src kinase using the string method with swarms-of-trajectories. Biophys. J. 97, L8–L10 (2009).

25. Yang, S., Banavali, N. K. & Roux, B. Mapping the conformational transition in Src activation by cumulating the information from multiple molecular dynamics trajectories. Proc. Natl. Acad. Sci. U. S. A. 106, 3776–3781 (2009).

26. Shirts, M. & Pande, V. S. Screen Savers of the World Unite! *Science* 290, 1903–1904 (2000).

27. Chopra, N. et al. Dynamic Allostery Mediated by a Conserved Tryptophan in the Tec Family Kinases. PLoS Comput. Biol. 12, e1004826 (2016).

28. Pande, V. S., Beauchamp, K. & Bowman, G. R. Everything you wanted to know about Markov State Models but were afraid to ask. Methods 52, 99–105 (2010).

29. Bowman, G. R., Pande, V. S. & Noé, F. An Introduction to Markov State Models and Their Application to Long Timescale Molecular Simulation. 797, (2014).

30. Nagar, B. et al. Structural basis for the autoinhibition of c-Abl tyrosine kinase. Cell 112, 859–71 (2003).

31. Pendergast, A. M. The Abl family kinases: Mechanisms of regulation and signaling. Advances in Cancer Research 85, 51–100 (2002).

32. Schwantes, C. R. & Pande, V. S. Improvements in Markov State Model Construction Reveal Many Non-Native Interactions in the Folding of NTL9. J. Chem. Theory Comput. 9, 2000–2009 (2013).

33. Noé, F. & Clementi, C. Kinetic distance and kinetic maps from molecular dynamics simulation. J. Chem. Theory Comput. 11, 5002–5011 (2015).

34. McGibbon, R. T. & Pande, V. S. Identification of simple reaction coordinates from complex dynamics. 1–16 (2016).

35. Pérez-Hernández, G. et al. Identification of slow molecular order parameters for Markov model construction. J. Chem. Phys. 139, 15102 (2013).

36. McGibbon, R. T. & Pande, V. S. Variational cross-validation of slow dynamical modes in molecular kinetics. J. Chem. phyics 142, (2015).

37. Nüske, F., Keller, B. G., Pérez-Hernández, G., Mey, A. S. J. S. & Noé, F. Variational Approach to Molecular Kinetics. J. Chem. Theory Comput. 10, 1739–52 (2014).

38. Xiao, Y., Liddle, J. C., Pardi, A. & Ahn, N. G. Dynamics of protein kinases: Insights from nuclear magnetic resonance. Acc. Chem. Res. 48, 1106–1114 (2015).

39. Xiao, Y. et al. Phosphorylation releases constraints to domain motion in ERK2. Proc. Natl. Acad. Sci. U. S. A. 111, 2506–11 (2014).

40. Gordon, J. C. et al. H++: a server for estimating pKas and adding missing hydrogens to macromolecules. Nucleic Acids Res. 33, W368–71 (2005).

41. Meng, Y., Lin, Y. L. & Roux, B. Computational study of the ‘DFG-Flip’ conformational transition in c-Abl and c-Src tyrosine kinases. J. Phys. Chem. B 119, 1443–1456 (2015).

42. Hinrichs, N. S. & Pande, V. S. Calculation of the distribution of eigenvalues and eigenvectors in Markovian state models for molecular dynamics. J. Chem. Phys. 126, 244101 (2007).

43. Noé, F. & Fischer, S. Transition networks for modeling the kinetics of conformational change in macromolecules. Curr. Opin. Struct. Biol. 18, 154–62 (2008).

44. Kornev, A. P., Haste, N. M., Taylor, S. S. & Eyck, L. F. T. Surface comparison of active and inactive protein kinases identifies a conserved activation mechanism. Proc. Natl. Acad. Sci. U. S. A. 103, 17783–8 (2006).

45. Cowan-Jacob, S. W. et al. The crystal structure of a c-Src complex in an active conformation suggests possible steps in c-Src activation. Structure 13, 861–71 (2005).

46. Johnson, L. N., Noble, M. E. M. & Owen, D. J. Active and inactive protein kinases: Structural basis for regulation. Cell 85, 149–158 (1996).

47. Yang, S. & Roux, B. Src kinase conformational activation: thermodynamics, pathways, and mechanisms. PLoS Comput. Biol. 4, e1000047 (2008).

48. Van Der Kamp, M. W. & Mulholland, A. J. Combined quantum mechanics/molecular mechanics (QM/MM) methods in computational enzymology. Biochemistry 52, 2708–2728 (2013).

49. Warshel, A. & Florian, J. in Encyclopedia of Computational Chemistry (John Wiley & Sons, Ltd, 2002). doi:10.1002/0470845015.cu0002

50. Berman, H. M. The Protein Data Bank. Nucleic Acids Res. 28, 235–242 (2000).

51. Eswar, N. et al. Comparative protein structure modeling using MODELLER. Curr. Protoc. Protein Sci. Chapter 2, Unit 2.9 (2007).

52. Salomon-Ferrer, R., Case, D. A. & Walker, R. C. An overview of the Amber biomolecular simulation package. Wiley Interdiscip. Rev. Comput. Mol. Sci. 3, 198–210 (2013).

53. Götz, A. W. et al. Routine Microsecond Molecular Dynamics Simulations with AMBER on GPUs. 1. Generalized Born. J. Chem. Theory Comput. 8, 1542–1555 (2012).

54. Salomon-Ferrer, R., Götz, A. W., Poole, D., Le Grand, S. & Walker, R. C. Routine Microsecond Molecular Dynamics Simulations with AMBER on GPUs. 2. Explicit Solvent Particle Mesh Ewald. J. Chem. Theory Comput. 9, 3878–3888 (2013).

55. Lindorff-Larsen, K. et al. Improved side-chain torsion potentials for the Amber ff99SB protein force field. Proteins 78, 1950–8 (2010).

56. Jorgensen, W. L., Chandrasekhar, J., Madura, J. D., Impey, R. W. & Klein, M. L. Comparison of simple potential functions for simulating liquid water. J. Chem. Phys. 79, 926 (1983).

57. Eastman, P. et al. OpenMM 4: A Reusable, Extensible, Hardware Independent Library for High Performance Molecular Simulation. J. Chem. Theory Comput. 9, 461–469 (2013).

58. Kohlhoff, K. J. et al. Cloud-based simulations on Google Exacycle reveal ligand modulation of GPCR activation pathways. Nat. Chem. 6, 15–21 (2014).

59. Sultan, M. M., Kiss, G., Shukla, D. & Pande, V. S. Automatic Selection of Order Parameters in the Analysis of Large Scale Molecular Dynamics Simulations. J. Chem. Theory Comput. 10, 5217–5223 (2014).

60. Mcgibbon, R. T. et al. MDTraj: a modern, open library for the analysis of molecular dynamics trajectories. bioRxiv 9–10 (2014).

61. Beauchamp, K. A. et al. MSMBuilder2: modeling conformational dynamics on the picosecond to millisecond scale. J. Chem. Theory Comput. 7, 3412–3419 (2011).

62. Pérez, F. & Granger, B. E. IPython: a System for Interactive Scientific Computing. Comput. Sci. Eng. 9, 21–29 (2007).

63. Hunter, J. D. Matplotlib: A 2D graphic environment. Comput. Sci. Eng. 9, 90–95 (2007).

64. Pedregosa, F. & Varoquaux, G. Scikit-learn: Machine learning in Python. J. Mach. Learn. 12, 2825–2830 (2011).

65. Humphrey, W., Dalke, A. & Schulten, K. VMD: visual molecular dynamics. J. Mol. Graph. 14, 33-8, 27-8 (1996).

66. Amitabh Varshney, Frederick P. Brooks, Jr., Jr. William, W. V. W. Linearly Scalable Computation of Smooth Molecular Surfaces. IEEE Comput. Graph. Appl. 14, (1994).

67. Frishman, D. & Argos, P. Knowledge-based protein secondary structure assignment. Proteins 23, 566–79 (1995).

